# A Method for Visual Psychophysics based on the Navigational Behaviour of Desert Ants (*Melophorus bagoti*)

**DOI:** 10.1101/2024.09.19.613987

**Authors:** Sudhakar Deeti, Vito Lionetti, Ken Cheng

## Abstract

The Australian desert ant *Melophorus bagoti* is known to navigate in their complex visual environment relying on path integration and landmark learning. We investigated the navigational behaviour of desert ants in response to visual stimulus changes along their route to a food source. Ants were trained to self-navigate a track between their nest and a feeder, with visual stimuli introduced near the feeder. After three days of training, we altered the visual scene by changing colours and observed the ants’ reactions. The results showed that ants in the test conditions, when visual changes were introduced, displayed more meandering paths, increased scanning behaviour, and slower speeds compared to control conditions. The sinuosity of their paths increased, and their orientation became less efficient. Additionally, changes in colour led to significant alterations in path characteristics, including a higher frequency of head oscillations and deviations from straight-line paths. These findings suggest that disruption to visual cues used for navigation causes noticeable changes in their movement patterns. A subset of these changes, easy to measure and calculate, can provide a signature to indicate that ants have detected a change in the visual stimuli. This proof-of-concept study thus highlights a method for studying visual psychophysics in ants in their natural habitat.

## INTRODUCTION

Psychophysics is a discipline that originated in the 19th century with scientists such as Ernst Weber, Gustav Fechner, Hermann von Helmholtz, Wilhelm Wundt, and Ernst Mach (Staley, 2021). A basic goal of the discipline is to map physical dimensions onto perceptual, psychological, or behavioural dimensions. An example of an aim might be estimating the magnitude of one kind of stimulus that is necessary for an animal to detect the stimulus, stimuli that are sensed by different sensory/perceptual systems such as lights, sounds, other mechanosensory stimuli, or chemicals (Pelli and Farell, 1995). Another aim might be to elucidate how an animal discriminates between different kinds of stimuli. Methodologically, psychophysical research requires test subjects to make judgements or produce a stimulus to match another stimulus. The research is concentrated on adult humans, and with these test participants, some instructions and a few practice trials would suffice before data collection could begin. The training time before psychophysical data could be collected is short. Other than adult humans, however, including non-verbal infants and other species (henceforth, “animals”), substantial training may be needed before psychophysical data collection could begin.

The required training exacts a cost in research, even though sometimes, the training could be short (Czaczkes and Kumar, 2020). To train laboratory pigeons in discrimination of visual textures, for example, took some 90 sessions in an operant chamber (Cook et al., 1995). Besides costs in time, sometimes, an animal may not be able to learn the operant task required. The well-studied North African desert ants *Cataglyphis fortis*, for example, could not be trained to discriminate black from white in a two-choice task, at least in the time that the investigators were willing to expend on the training (Schwarz and Cheng, 2010). For these reasons, it would be beneficial to do psychophysical experiments with no or minimal training required.

These minimal-training tasks typically examine the reaction of animals to stimuli. Perhaps at the minimal end is a method sending flies, *Drosophila melanogaster*, through an automated apparatus and recording their behaviours (Evans et al., 2011). As the flies walked through the maze, different kinds of visual stimuli were presented. Different populations, such as different genetically manipulated strains, could be compared in their reactions. Another kind of method relies on reactions to novel stimuli. Animals are presented one kind of stimulus repeatedly, a habituating stimulus. On a crucial test, they are presented a physically different stimulus. If the animal can detect a difference perceptually or psychologically, then it should show some reaction. If it cannot, a similar response as the previous presentation of the habituating stimulus should take place. This methodology is sometimes termed, loosely, “habituation/dishabituation”, although this characterisation is technically incorrect. Dishabituation refers to the reaction to the habituating stimulus after some novel stimulus has been presented, not the reaction to the novel stimulus (Rankin and Carew, 1988).

This reaction-to-novelty method can be and has been conducted on infants. Infants presented repeatedly pictures showing two objects, different objects on different trials, soon looked less and less, the familiar process of habituation taking place (Starkey et al., 1990). This ‘training’, if it should be called that at all, does not require the infants to master any task. When subsequently shown a picture with three objects, their looking increased, compared with a further presentation of a picture with two objects. The results suggested to the authors that 6- to 8-month-old infants possessed some concept of numerosity. Our goal in this account is to test such a method on desert ants as a method for conducting future visual psychophysical research, in addition to characterising in detail desert ants’ reactions to novel visual stimuli in the context of navigation.

Ants are said to display advanced cognition (Czaczkes, 2022), considered as cognition that “usually requires symbolic computation, abstract representation of information, mental model building, or information processing in the absence of current exposure to relevant external stimuli” (p. 51). Their navigational prowess is legion and has been well studied (Wehner, 2020; Freas and Cheng, 2022; Rössler, 2023). A recent focus features oscillatory behaviours of ants as they navigate (Murray et al., 2020; Clement et al., 2023). Our recent observations on the study species of this article, the Australian red honey ant *Melophorus bagoti*, revealed that these ants oscillate both their heads and their paths as they navigate to a feeder to back home, even on a familiar route (work in preparation). They swing their heads regularly left and right, and the travelled path ‘wiggles’ left and right. When encountering some change in their visual scenery, the oscillations change. Therein lies the route to a visual psychophyical research program.

We set up a track for the ants and provisioned them with food in a feeder near one end of the track, the other end encircling their nest. No explicit training is needed to train the desert ants to find the feeder and take food home: they ‘train themselves’ to find the feeder and go home with food. Visual stimuli were presented just before they reached the feeder. After sufficient self-training on the outbound route, we could make make an obvious change in the stimuli and note the ants’ reactions. Our goal in the proof of concept is not to do any visual psychophysics, but to find the best and easiest suite of measures to use as a signature that the ants had noticed a difference. In the process, because we must examine many measures to come up with the most diagnostic suite, we also examine the suite of changes that come with visual scene changes on the route of this desert ant. We explain in the Discussion how such a method can be used for programs of research in visual psychophysics conducted in the field, in the natural habitat of this and potentially other ant species.

## METHODS

### Study species

In this study, we used a salient change of colour as a way to elicit change in the navigational behaviours of foragers. We conducted the tests on a single *Melophorus bagoti* nest (Lubbock, 1883) during the period of November 2023 and February 2024. The nest was situated close to the Centre for Appropriate Technology, located 10 km south of Alice Springs, NT, Australia (23°45′28.12″S, 133°52′59.77″E). The nest’s surroundings comprised an expansive open space within a semi-arid desert landscape, dotted with buffel grass (*Pennisetum cenchroides*), a mix of *Acacia* bushes, and *Eucalyptus* trees (Deeti and Cheng 2021a; Deeti and Cheng 2021b; Deeti et al., 2020; Deeti et al., 2023a), which formed a distinct visual panorama. In Australia, there are no specific ethical regulations pertaining to the study of ants, and our experimental procedures were entirely non-invasive.

The red honey ant, *M. bagoti*, is the most thermophilic ant species found on the Australian continent (Christian and Morton, 1992). Our observations revealed that these ants were usually active outside the nest when temperatures ranged between 32 and 40°C. During the hot southern summer, these *M. bagoti* ants engage in foraging. They scavenge primarily dead spiders, sugar ants, termites, centipedes, moths and other dead arthropods. Additionally, they also collect sugary plant exudates and seeds (Muser et al., 2005; Schultheiss and Nooten, 2013). The *M. bagoti* foragers typically forage solitarily, covering distances up to 50 m from the nest (personal observations), relying on path integration and the terrestrial visual surround rather than chemical trails to navigate (Cheng et al., 2009).

### Set up

Before commencing the experiment, the grass in the area between the nest and the feeder was cleared, while the surrounding area remained populated with bushes and trees. The foragers were trained to follow a straight outbound route to a feeder located approximately 7 m from the nest, which contained a mix of Arnott brand cookie pieces and mealworms to attract foraging ants. To guide the ants towards the feeder, a plywood barrier around the route was constructed. This barrier spanned 10 m in length and 1.4 m in width. It was built using PVC floor tiles, each measuring 90 cm in length and 15 cm in height, buried 5 cm deep into the ground for stability. The PVC floor tiles created a slippery surface for ants, preventing them from climbing and diverting from the designated foraging path. At a distance of 6 m from the nest, two symmetrical plywood boards, each measuring 180 × 120 cm, were positioned on the ants’ foraging route, with a 50-cm gap between them. This gap intersected a line connecting the nest, the centre of the recording area, and the feeder. Before the plywood blocks, fine white sand was spread on the ground within the recording area to ensure clear visibility of the red ants against the sandy red soil background. This ensured a high contrast between the ants and the background, making them easily distinguishable in recorded footage. No notable difference was observed in the behaviour of the ants on white sand versus red soil.

The trajectories of foragers when going to collect the food from the feeder (outbound) were recorded in the area before the boards. To capture the ant paths, a tripod was set up at a height of 1.2 m from the ground. A Sony Handy camera (FDR-AX700) was mounted on the tripod, and the recording was done at a frame rate of 25 frames per second (fps). The recording area had a size of 1 m by 1 m, and the camera had a resolution of 3860 by 2160 pixels.

### Experimental procedure

In this experiment, we utilized three distinct groups of painted ants from the nest, each assigned to one of three colour-change conditions. The conditions differed in the colours before and after a colour change imposed in the course of the experiment: black to white (n=20, ants painted yellow), white to black (n=20, ants painted white), white to orange (n=20, ants painted pink). The aim was to examine how altering colours in their environment affects the navigational behaviour of *Melophorus bagoti* ants, in order to find a suite of measures indicative of ants’ detecting a visual change. In the black–white condition, we initially trained the ants to recognize and habituate to black boards around the gap leading to the feeder. Newly arriving unpainted ants at the feeder were painted yellow. Subsequently, over three days, the painted ants were observed and allowed to learn the route. On the third day, the first appearance on an outbound trip by each ant was recorded as a ‘before’ measure. Each recorded ant was marked with an extra dot of colour for individual identification. On the fourth day, black boards were replaced with white ones before the ants were active, and individual foragers’ behaviour was recorded on their first visit for the ‘after’ measure. Upon reaching the feeder, ants were marked with a black dot to prevent double testing. After completing the ‘after’ test, ants were excluded from further study, that is, ignored while they continued to go about their foraging. This process was repeated for the other groups undergoing changes of colour from white to black and from white to orange, with each group trained and tested in the same fashion except for the colours of the boards.

### Tracking and Data extraction

We used the animal tracking program DLTdv8 (version 8.2.9) in MATLAB (2022B) to extract frame-by-frame coordinates of the foragers’ head and thorax—the tip of the head and the middle of the thorax (see Figure S1). During the tests, ants frequently displayed a series of stereotypical successive fixations (<0.1 mm/s) in different directions by rotating on the spot at one location, known as a “scanning bout” and we extracted the number and duration of scanning bouts (Deeti et al. 2023b).

For orientations, each change in direction of heading from left to right or from right to left was taken as an inflection point, again with a minimum sequence length of 3 frames.

Temporal frequencies and amplitudes, the latter measured as degrees of orientation change, could then be calculated in an analogous fashion to the calculations for path-oscillatory characteristics. Spatial frequencies do not make any sense in the context of head orientations, as position in space does not form a defining role in the thorax–head direction.

### Data Analysis

We used the extracted frame-by-frame coordinates to calculate the final heading direction, speed, orientation direction, orientation angular velocity and path characteristics. We determined the final heading direction by the head–thorax direction as an ant exited the recording area. This vector represented the orientation of each ant at that specific moment. To understand how quickly and directly the ants moved at each displacement location, we calculated the speed of the workers along with several characteristics of the trajectory. These path characteristics were based on the positions of the thorax across frames. For each ant, speed refers to the magnitude of an ant’s velocity and was calculated as path distance covered by the entire trajectory divided by the duration of the trip, excluding the stopping durations. We measured the speed at which ants change their orientation direction by observing how quickly their orientation changes over time. As ants walk, they turn their heads left and right in oscillation. The orientation direction of each frame was defined by the thorax–head vector, so that the orientation angular velocity is measured as the change in this direction per unit time.

For path characteristics on recorded runs, we used three indices of straightness: *path straightness, sinuosity*, and 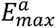, each of which relates to the directness of navigation towards a destination. *Straightness* of the trajectory is computed as the ratio of the straight-line distance (*D*) traversed by the recorded trajectory to the overall length (*L*) of the path (Batschelet, 1981; Deeti et al., 2023c; Islam et al., 2021; Islam et al., 2023). Therefore, the formula for straightness is typically expressed as *D/L*. *Straightness* ranges from 0 to 1, with larger values indicating straighter paths, while smaller values indicate more curved or convoluted paths. *Sinuosity* is an estimate of the tortuosity of the recorded path, calculated as 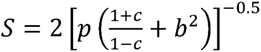, where *p* is the mean step length, *c* is the mean cosine of turning angles and *b* is the coefficient of variation of the step length. A trajectory *step* is the movement of the animal’s thorax position recorded in consecutive video frames. Accordingly, step lengths are the Euclidean distances between consecutive points along a path, and turning angle refers to the change in direction between two consecutive steps. Sinuosity varies between 0 (straight) and 1 (extremely curved) (Benhamou, 2004; Lionetti et al., 2023). The maximum expected displacement of a path, 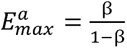, where β is the mean cosine of turning angles, is a dimensionless value expressed as a function of the number of steps and is consistent with the intuitive meaning of straightness (Cheung et al. 2007). Larger maximum expected displacement values indicate straighter paths with a greater displacement in the trajectory, while a smaller value suggests more localised or constrained movement. Paths were characterized and visualized in R (version 4.2.1; R Core Team, 2020) using the packages trajr (McLean and Skowron Volponi 2018) and Durga (Khan and McLean 2023).

Ants frequently displayed a series of successive fixations in different head directions by rotating on the spot at one location, known as a “scanning bout” (Deeti, 2023b; Islam et al., 2022). For each ant in each condition, we extracted the number of scanning bouts and the scanning-bout durations (from the start of a scanning bout until the ant started walking again).

To characterize oscillations of paths, we calculated amplitude, temporal frequency, and spatial frequency. Initially, we segmented ‘wiggles’ by identifying inflection points on the path, at which the path changed from turning right to turning left or vice versa. The amplitude of each wiggle was the point with the maximum distance from the straight line connecting the end points of each segment, tallied in centimetres. The spatial period of the wiggle was calculated from the length of the straight line between the inflection points defining a wiggle and doubling it (one wiggle being half a cycle), also in centimetres. The spatial frequency was then derived by taking the inverse of the spatial period (cycles/cm). To calculate the temporal period of a wiggle, we counted frames between inflection points and doubled the count. The number of frame changes was translated into seconds by dividing by the frame rate of 25 f/s. The temporal frequency was the inverse of the temporal period (cycles/s). We excluded the first and last segments due to unknown inflection points beyond the data range. For each of the remaining segments, we calculated the average and standard deviation across segments for frequencies and amplitudes, plotting individual data points across ants for visualization. For all these measures, we also calculated the within-individual standard deviation (SD) across all the segments (wiggles) of each ant’s path.

For head orientations, each change in direction of heading from left to right or from right to left was taken as an inflection point, with a minimum sequence length of 3 frames as a requirement. Temporal frequencies and amplitudes, the latter measured as degrees of orientation change, could then be calculated in an analogous fashion to the calculations for path-oscillatory characteristics. Spatial frequencies do not make any sense in the context of head orientations, as position in space does not form a defining role in the thorax–head direction.

With two separate oscillations, a third parameter can be calculated, that of the phase relationship between head-orientation and path oscillations. Head orientation changed faster than the path ‘wiggle’, so that we plotted the former on the latter cycle. Double wiggles of the path were represented as a circle with 0° representing the start of the turn to the left. Angles thus represented proportions of a period. We plotted the start of head-orientation cycles (start of turn to the left) on this circle, increasing clockwise, combining all ants and all segments. We repeated this analysis defining 0° as the start of turn to the right. Because the wiggles are slightly unequal in duration, if the start of the left turn is 0°, the start of the right turn does not fall in general exactly at 180°, but only somewhere in its vicinity.

We show scanning data as violin plots to better depict many data points bunched at the same level. We show oscillatory measures as individual data points across pairs of associated control and experimental conditions. Other measures are shown as box plots.

### Data Analysis

The experiments conducted with the two different colours in training and three different colours as tests were analysed using paired t-tests, one test for each control and its associated test colour regime. Statistical analyses were conducted using R (version 4.3.1). Alpha was set at 0.01 because the same data set was used to compute several dependent variables. We also measured the effect size of all dependent variables for identifying a group that could form an easy signature of detection of visual change (Table 1). As a measure of effect size, we used Cohen’s d, determined by calculating the mean difference between the control and test groups, and then dividing the result by the pooled standard deviation.

**Table 1:** The effect size of variables measured.

|  | Group | Effect size | Cohen's (*d*) Value |
| --- | --- | --- | --- |
| Sinuosity | BC Vs. BT | 0.3 | -4.7 |
|  | WC Vs. WT | 0.2 | -1.7 |
|  | OC Vs. OT | 0.2 | -2.5 |
| E<sup>a</sup><sub>Max</sub> | BC Vs. BT | -964.2 | 2.1 |
|  | WC Vs. WT | -446.8 | 1.6 |
|  | OC Vs. OT | -973.7 | 1.6 |
| Straightness | BC Vs. BT | -0.4 | 3.1 |
|  | WC Vs. WT | -0.3 | 1.7 |
|  | OC Vs. OT | -0.4 | 2.5 |
| Speed | BC Vs. BT | -180.6 | 4.7 |
|  | WC Vs. WT | -111.4 | 1.6 |
|  | OC Vs. OT | -153.6 | 2.4 |
| Orientation angular velocity | BC Vs. BT | -64.24 | 1.19 |
|  | WC Vs. WT | -53.7 | 1.2 |
|  | OC Vs. OT | -218.5 | 5.2 |
| Duration of Walk | BC Vs. BT | 13.71 | -1.4 |
|  | WC Vs. WT | 6.4 | -1.6 |
|  | OC Vs. OT | -2.6 | -4.4 |
| Number of Scans | BC Vs. BT | 3.2 | 1.4 |
|  | WC Vs. WT | 5.4 | 2.4 |
|  | OC Vs. OT | 12.7 | 1.1 |
| Scanning bout Duration | BC Vs. BT | 4.3 | 1.8 |
|  | WC Vs. WT | 56.9 | 1.7 |
|  | OC Vs. OT | 34.3 | 1.6 |
| Amplitude Mean (H) | BC Vs. BT | -4.8 | -1.09 |
|  | WC Vs. WT | -6.5 | -1.06 |
|  | OC Vs. OT | -4.3 | -0.7 |
| Amplitude SD (H) | BC Vs. BT | -7.5 | -1.1 |
|  | WC Vs. WT | -12.5 | -1.4 |
|  | OC Vs. OT | -9.2 | -1.08 |
| Temporal Frequency mean (H) | BC Vs. BT | -7.5 | -12.2 |
|  | WC Vs. WT | 0.4 | 1.09 |
|  | OC Vs. OT | 0.4 | 1.01 |
| Temporal Frequency SD (H) | BC Vs. BT | -7.5 | -17.4 |
|  | WC Vs. WT | -0.1 | -0.4 |
|  | OC Vs. OT | 0.4 | 1.02 |
| Amplitude Mean (T) | BC Vs. BT | -4.4 | -1.06 |
|  | WC Vs. WT | -6.7 | -0.5 |
|  | OC Vs. OT | 6.2 | 0.5 |
| Amplitude SD (T) | BC Vs. BT | -8.1 | -1.2 |
|  | WC Vs. WT | -13.2 | -0.8 |
|  | OC Vs. OT | 6.6 | 0.4 |
| Temporal Frequency mean (T) | BC Vs. BT | -8.1 | -13.3 |
|  | WC Vs. WT | 0.3 | 0.8 |
|  | OC Vs. OT | 2.08 | 4.6 |
| Temporal Frequency SD (T) | BC Vs. BT | -8.1 | -19.3 |
|  | WC Vs. WT | -0.1 | -0.4 |
|  | OC Vs. OT | 2.08 | 5.2 |
| Spatial Frequency mean (T) | BC Vs. BT | -0.01 | -1.1 |
|  | WC Vs. WT | -0.02 | -1.5 |
|  | OC Vs. OT | 0.008 | 0.6 |
| Spatial Frequency SD (T) | BC Vs. BT | -0.01 | -0.8 |
|  | WC Vs. WT | -0.03 | -1.7 |
|  | OC Vs. OT | 0.006 | 0.3 |

## RESULTS

We compared experienced foragers’ performance on the fourth morning when they faced a changed colour of the boards enroute to the feeder (the Test condition) with their performance on the third day, before any visual changes (the Control condition). On the morning of the third day, recorded paths showed that the foragers were feeder-oriented, exhibiting quite straight paths with little wiggle and meandering as well as few scans (Figure 2, Figure S2).

**Figure 1.**
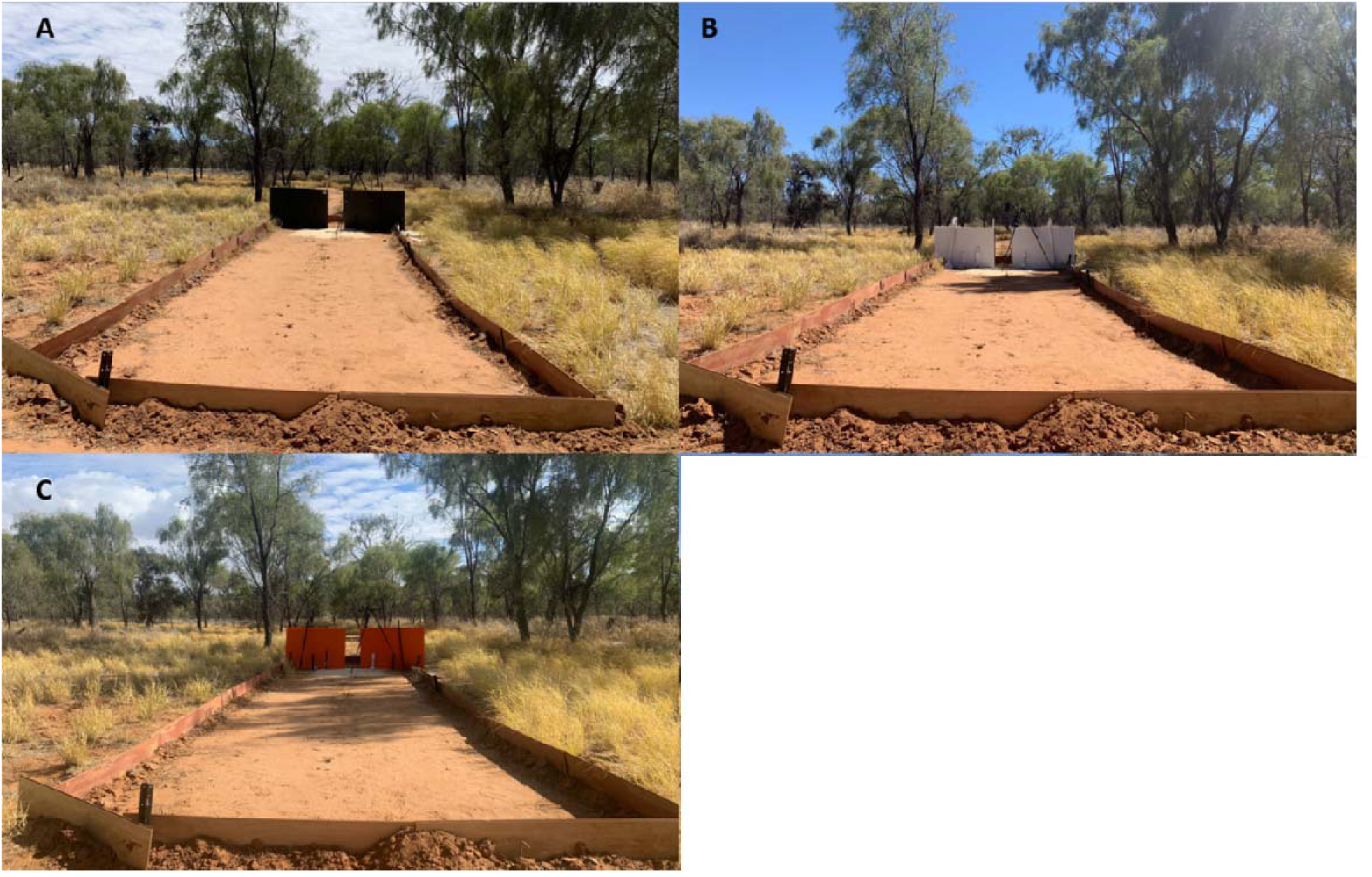
The different experimental colour boards set up at the recording site. (A) black boards. (B) white boards, (C) orange boards. Ants were trained with different colours before and after the colour change.

**Figure 2.**
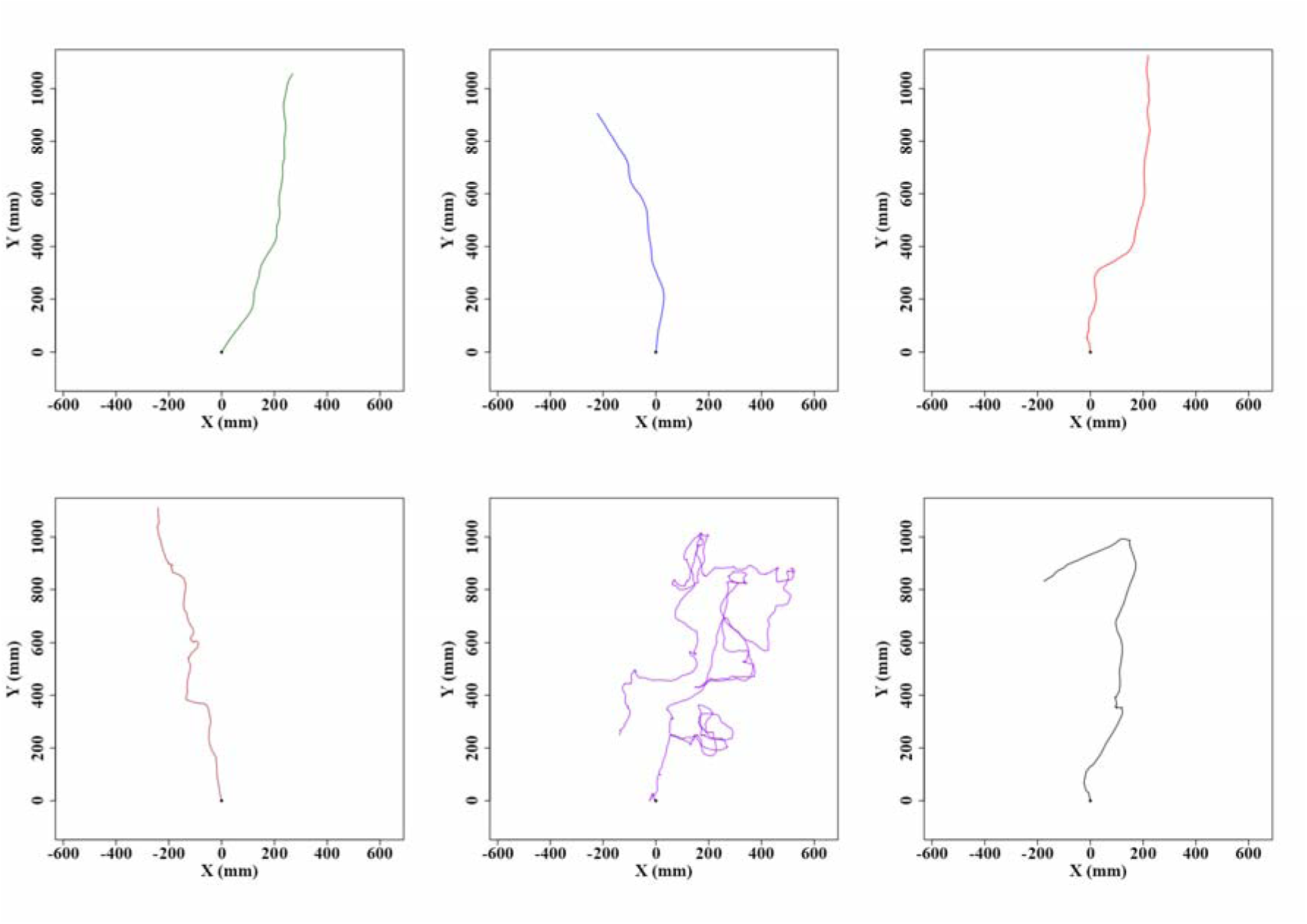
Trajectories of red honey ants in control and test conditions with various colours. (A) to (F) show the Control and Test paths of the individual desert ants during foraging. (A), (B), and (C) represent the control paths before the colour of the board changed. (D), (E) and (F) depict the test paths after the colour changes. (A), (D), Black to White; (B), (E), White to Orange; (C), (F), White to Black. Each trajectory plot represents the movement path of an individual ant. The y-axis points towards the goal. The coordinates (0,0) represent the starting frame of each recording.

These characteristics together suggest competent navigation. In the test conditions, foragers reacted to the panoramic colour change, with most ants meandering until they passed the gap and performing a greater number of scans along their route with much wiggling. In describing results in detail, we call each associated pair of conditions by the test colour; thus the white- to-black change was called “Black control” and “Black test”.

In path characteristics, ants exhibited a more curved and meandering trajectory in the colour- change conditions compared to the control conditions. We found differences across conditions in each of our three measures of path meander. Firstly, sinuosity increased significantly in each colour-change condition (Black control vs. Black test: *t* = –7.06, *df* = 23, *p* ≤ 0.0003; White control vs. White test: *t* = –11.1, *df* = 37.9, *p* ≤ 0.0001; Orange control vs. Orange test: *t* = –7.92, *df* = 35.2, *p* ≤ 0.0001) (Figure 3A). Secondly, 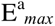 was lower in the colour-change conditions, with the ants having a smaller amount of displacement per unit length travelled compared to the control conditions. The t-tests showed a significant difference in test conditions compared to the control conditions in all comparisons (Black control vs. Black test: *t* = 6.4, *df* = 19.3, *p* ≤ 0.0001; White control vs. White test: *t* = 5.5, *df* = 19.1, *p* ≤ 0.0001; Orange control vs. Orange test: *t* = 4.6, *df* = 19.1, *p* = 0.0001) (Figure 3B). With straightness, the colour change led to a significantly higher magnitude of path deviation from a straight- line path compared to paths in the Control conditions, again in all comparisons (Black control vs. Black test: *t* = 6.2, *df* = 19.2, *p* ≤ 0.0005; White control vs. White test: *t* = 8.96, *df* = 21.2, *p* ≤ 0.0001; Orange control vs. Orange test: *t* = 7.98, *df* = 20.8, *p* ≤ 0.0001) (Figure 3C). Colour change had a noticeable impact on the speed and orientation angular velocity of the foragers. When the foragers faced a change in the colour of the stimuli, they slowed down, frequently shifted their gaze in different directions, and stayed in the recording area longer compared to the respective controls (Figure 3). In speed, the t-tests revealed significant differences in all comparisons (Figure 3D), (Black control vs. Black test: *t* = 8.2, *df* = 29.2, *p* ≤ 0.0004; White control vs. White test: *t* = 8.8, *df* = 30.9, *p* ≤ 0.0001; Orange control vs. Orange test: *t* = 7.8, *df* = 36.1, *p* ≤ 0.0001). Changes in odour also decreased the magnitude of orientation angular velocity in the foragers (Figure 3E). The t-tests showed significant differences between two pairs in orientation angular velocity (Black control vs. Black test: *t* = 3.6, *df* = 35.1, *p* ≤ 0.0009; White control vs. White test: *t* = 3.9, *df* = 31.04, *p* = 0.0004) but not with Orange control vs. Orange test: *t* = –0.19, *df* = 35.7, *p* = 0.847). Changes in colour also increased the duration of the foragers in the recording area (Figure 3F). The t-tests showed significant differences between each of the pairs in duration (Black control vs. Black test: *t* = –5.5, *df* = 19.2, *p* ≤ 0.0002; White control vs. White test: *t* = –5.1, *df* = 23.01, *p* ≤ 0.0001; Orange control vs. Orange test: *t* = –2.7, *df* = 28.6, *p* = 0.009).

**Figure 3.**
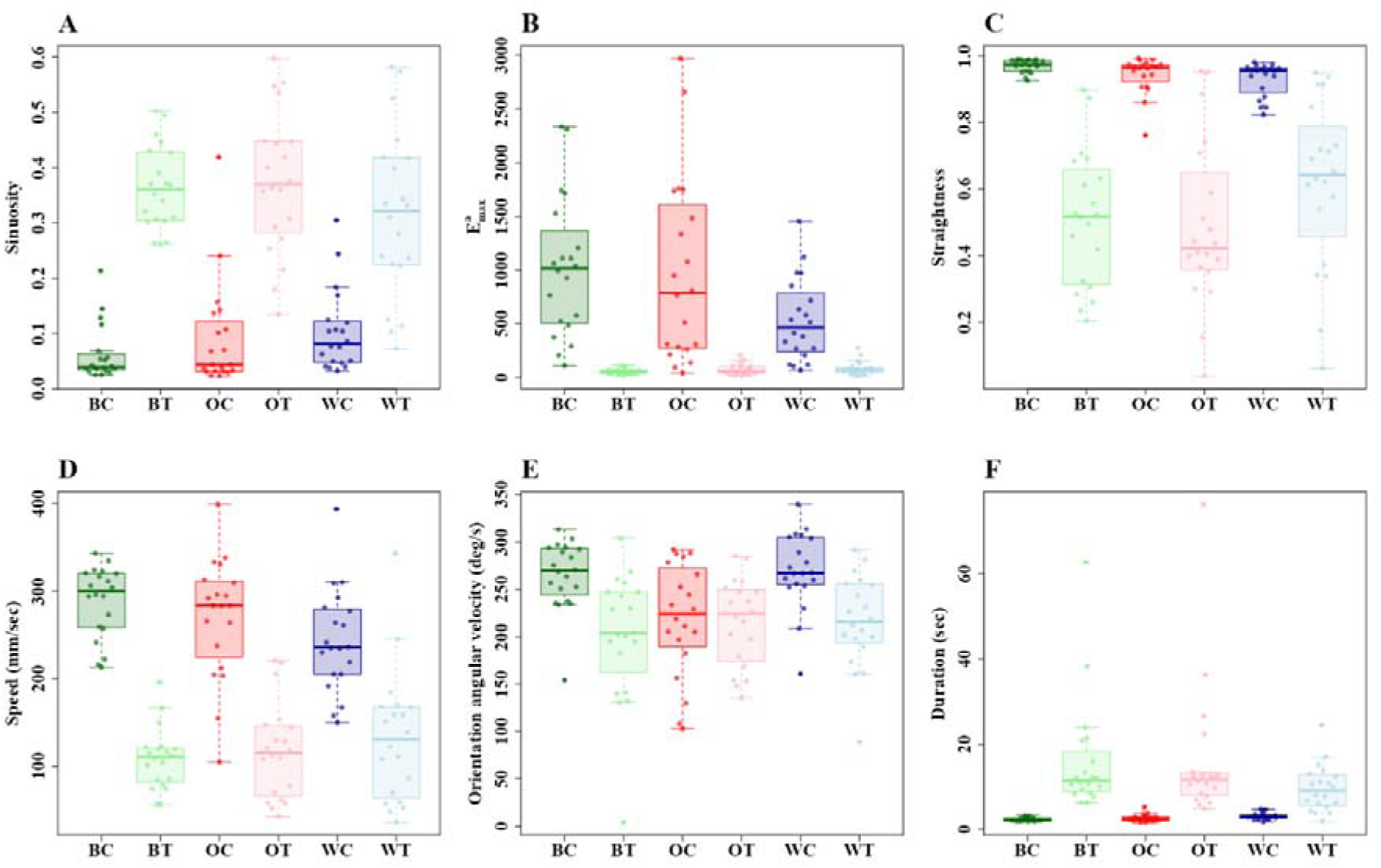
Path characteristics of red honey ants during the control and test runs. Shown are the (A) Sinuosity, (B) E^a^_max_, (C) Straightness, (D) Mean speed, (E) Orientation angular velocity and (F) duration of walk in the recording area under control and test observations. Box plots represent the median (line inside the box), interquartile range (box), and extreme values excluding outliers (whiskers). Individual data points are shown as dots. Each point represents a single trajectory measure. BC, WC and OC represent Black control, White control, Orange control, whereas BT, WT and OT represent Black, White, Orange tests, respectively.

Colour change increased the scan number substantially in foragers. In Control recordings, i.e. those without any colour change, we see few scans by the foragers (Figure 4A); most ants did not perform any scans at all in the recording area and the average number of scans was 0.2 per metre. In contrast, in the colour-change conditions, every forager scanned at least twice, and the average number of scans was 3.4 per metre (Figure 4A). This reflects a significant increase of scans in the Test conditions when compared with the controls in all comparisons (Black control vs. Black test: *t* = –4.5, *df* = 19.6, *p* ≤ 0.0001; White control vs. White test: *t* = –7.6, *df* = 20.3, *p* ≤ 0.0007; Orange control vs. Orange test: t = –3.5, df = 19.04, *p* = 0.002). The durations of scanning bouts also increased significantly in the test conditions. In durations, the t-tests revealed significant differences in all comparisons (Figure 4B) (Black control vs. Black test: *t* = –6.5, *df* = 20.2, *p* = 0.00005; White control vs. White test: *t* = –5.1, *df* = 22.4, *p* ≤ 0.0001; Orange control vs. Orange test: *t* = –3.7, *df* = 29.2, *p* = 0.008).

**Figure 4.**
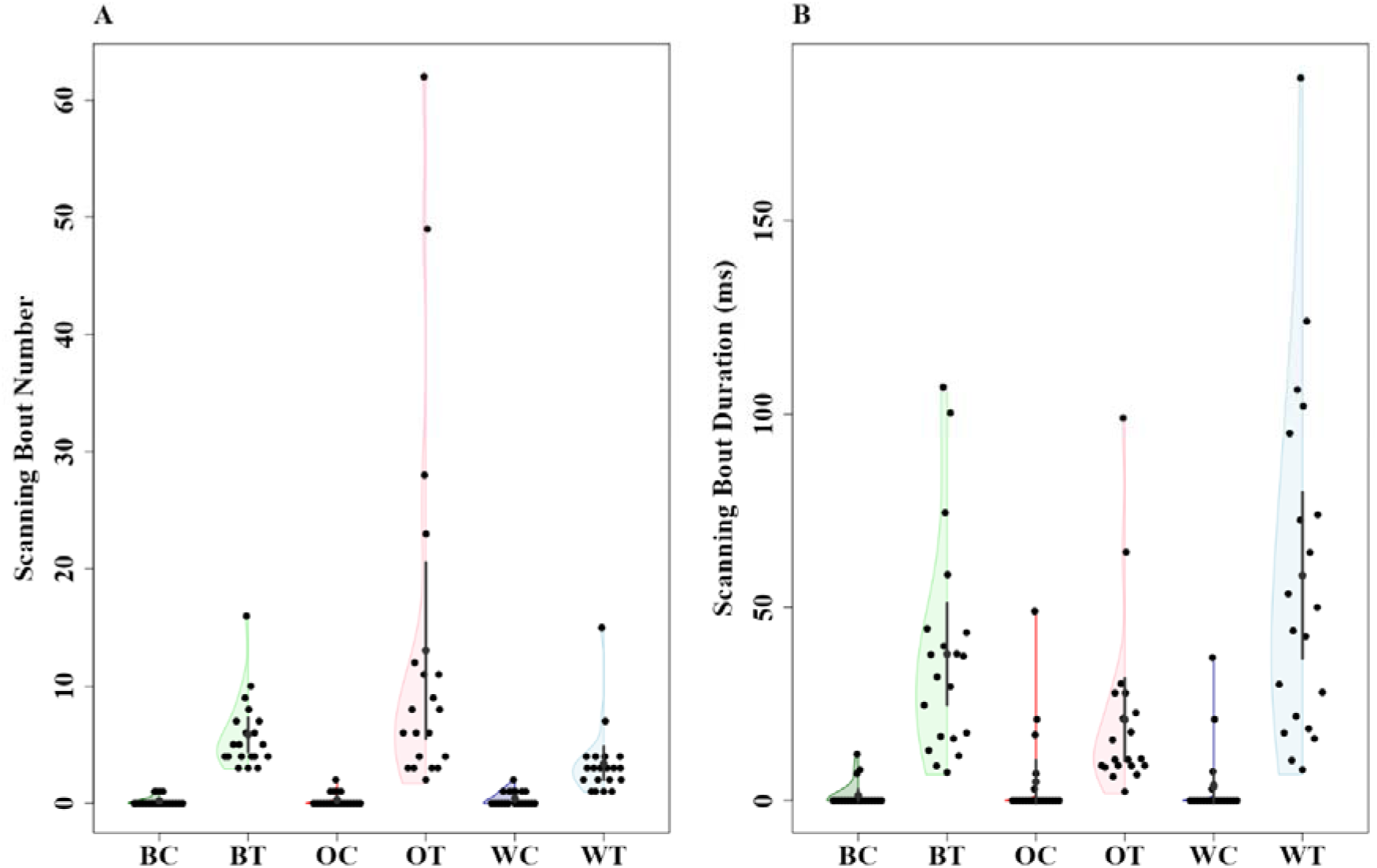
Number of scanning bouts and the duration of scanning bouts of the red honey ants during the control and test observations. The half-violin plots in (A) and (B) show the distribution of the mean number of scanning bouts and duration of the scanning bouts, respectively, of ants across control and test trials. In the half-violin plots, the solid dot shows the mean while the vertical bar represents the 95% confidence interval of the mean. BC, WC and OC represent Black control, White control, Orange control, whereas BT, WT and OT represent Black, White, Orange tests, respectively.

Ants oscillated their heads laterally as they walked. We analysed the amplitude and temporal frequency of these oscillations in the experimental test and control conditions. In the colour- change conditions, orientation-oscillation amplitude mean and SD were higher compared to controls. Together with slower forward movement with a colour change, ants also showed slower head movements resulting in a lower temporal frequency of orientation oscillations. The paired t-tests showed significant differences in Amplitude mean in all comparisons (Figure 5A) (Black control vs. Black test: *t* = –3.5, *df* = 22.1, *p* = 0.001; White control vs. White test: *t* = –2.9, *df* = 33.06, *p* = 0.005; Orange control vs. Orange test: *t* = –2.7, *df* = 34.5, *p* = 0.009). Amplitude SDs also differed significantly in all comparisons (Figure 5B) (Black control vs. Black test: *t* = –3.7, *df* = 21.1, *p* = 0.001; White control vs. White test: *t* = –4.1, *df* = 31.1, *p* = 0.0002; Orange control vs. Orange test: *t* = –3.7, *df* = 30.3, *p* = 0.007). The paired t-tests showed significant differences in Temporal frequency in all comparisons (Figure 5C), (Black control vs. Black test: *t* = 3.2, *df* = 24.2, *p* = 0.003; White control vs. White test: *t* = 5.54, *df* = 22.06, *p* = 0.00001; Orange control vs. Orange test *t* = 3.9, *df* = 33.2, *p* = 0.0003). The SD of temporal frequency not differed significantly in all comparisons (Black control vs. Black test: *t* = 2.8, *df* = 19.9, *p* = 0.01 White control vs. White test: *t* = 1.6, *df* = 24.5, *p* = 0.09; Orange control vs. Orange test: *t* = 3.5, *df* = 27.5, *p* = 0.001) (Figure 5D).

**Figure 5.**
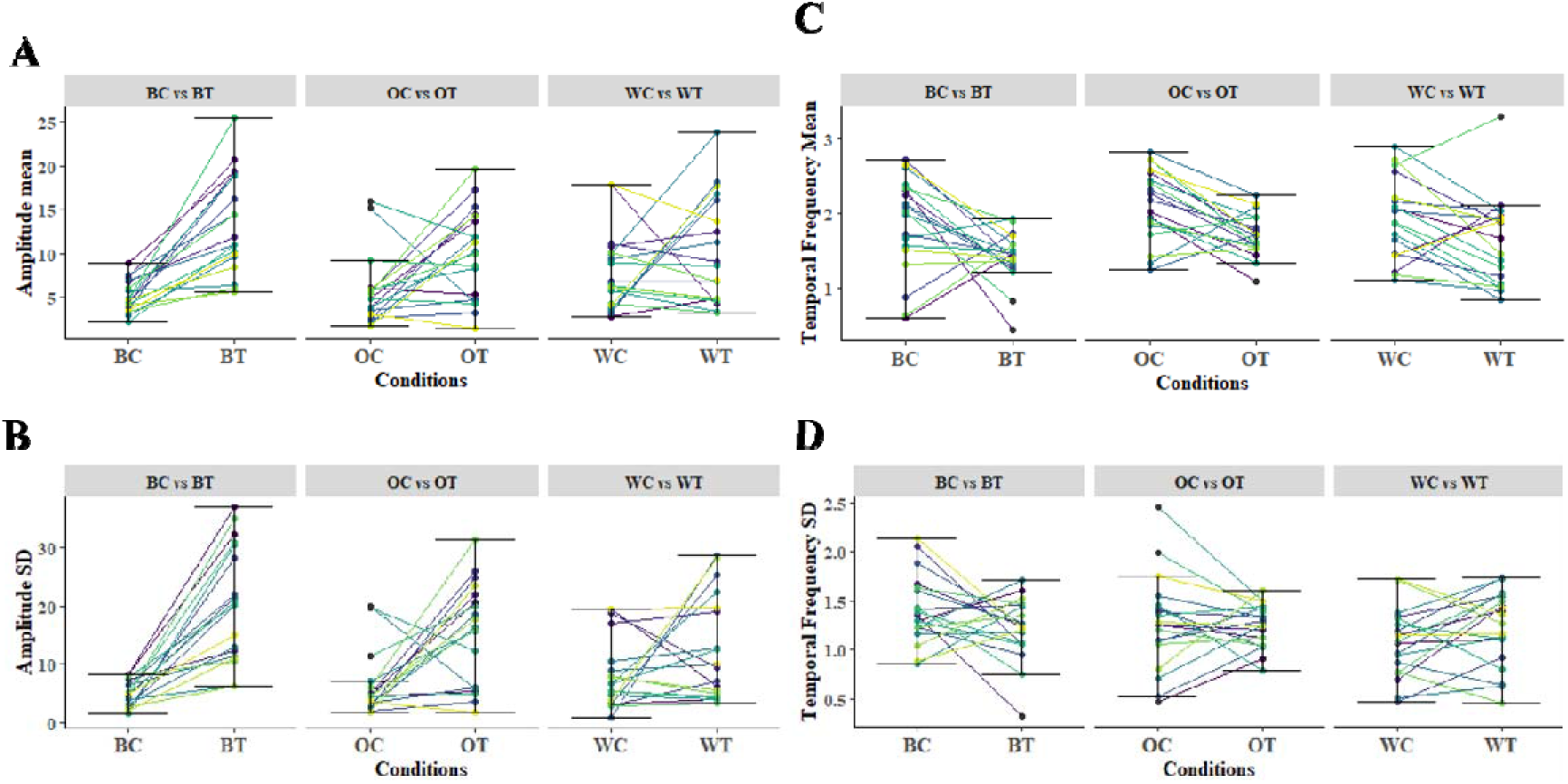
Head orientation oscillations. The plots show the effects of colour change on Amplitude mean, Temporal frequency mean, Amplitude SD and Temporal frequency SD from the control to the test conditions. BC, WC and OC represent Black control, White control, Orange control, whereas BT, WT and OT represent Black, White, Orange tests, respectively. Units are degrees for amplitude and cycles per second for temporal frequency.

Then we examined the effect of colour change on path oscillations, examining means and SDs in Amplitude, Temporal frequency, and Spatial frequency. Spatial frequency was affected by colour change, with no consistent pattern for temporal frequency or amplitude. The paired t-test on Amplitude showed significant differences in two comparisons (Black control vs. Black test: *t* = –3.4, *df* = 30.4, *p* = 0.001; White control vs. White test *t* = –2.6, *df* = 33.3, *p* = 0.01), but not with or Orange control vs. Orange test (*t* = –1.7, *df* = 21.7, *p* = 0.09) (Figure 6A). Amplitude SD also showed inconsistent effects across comparisons in the t-tests (Black control vs. Block test: *t* = –4.07, df = 25.01, *p* = 0.0004; White control vs. White test: *t* = –2.6, *df* = 21.8, *p* = 0.014, not significant; Orange control vs. Orange test: *t* = –3.9, *df* = 31.2, *p* = 0.0004) (Figure 6B). In Spatial frequency mean, the paired t-tests showed consistent significant differences in all comparisons (Black control vs. Black test: *t* = –3.6, *df* = 20.8, *p* = 0.001; White control vs. White test: *t* = –5, *df* = 19.7, *p* ≤ 0.0001; Orange control vs. Orange test: *t* = –5.4, *df* = 22.3, *p* ≤ 0.0001) (Figure 6C). In Spatial frequency SD as well, the t-tests revealed significant differences in all comparisons (Black control vs. White test: *t* = –2.8, *df* = 19.6, *p* = 0.009; Black control vs. Black test: *t* = –5.5, *df* = 22, *p* ≤ 0.0001; Orange control vs. Orange test: *t* = –5.6, *df* = 20.1, *p* ≤ 0.0001) (Figure 6D). In Temporal frequency means, results were inconsistent across comparisons. The paired t-tests showed significant differences in Black control vs. Black test (*t* = 2.6, *df* = 33.9, *p* = 0.01), but not with White control vs. White test (*t* = 0.4, *df* = 37.3, *p* = 0.68) or Orange control vs. Orange test (*t* = 2.1, *df* = 36.2, *p* = 0.03) (Figure 6E). Temporal frequency SD, on the other hand, did not show significant differences in any comparison (Black control vs. Black test: *t* = 0.03, *df* = 35.7, *p* = 0.97; White control vs. White test: *t* = 1.2, *df* = 37.9, *p* = 0.21; Orange control vs. Orange test: *t* = 1.77, *df* = 36.1, *p* = 0.32) (Figure 6F).

**Figure 6.**
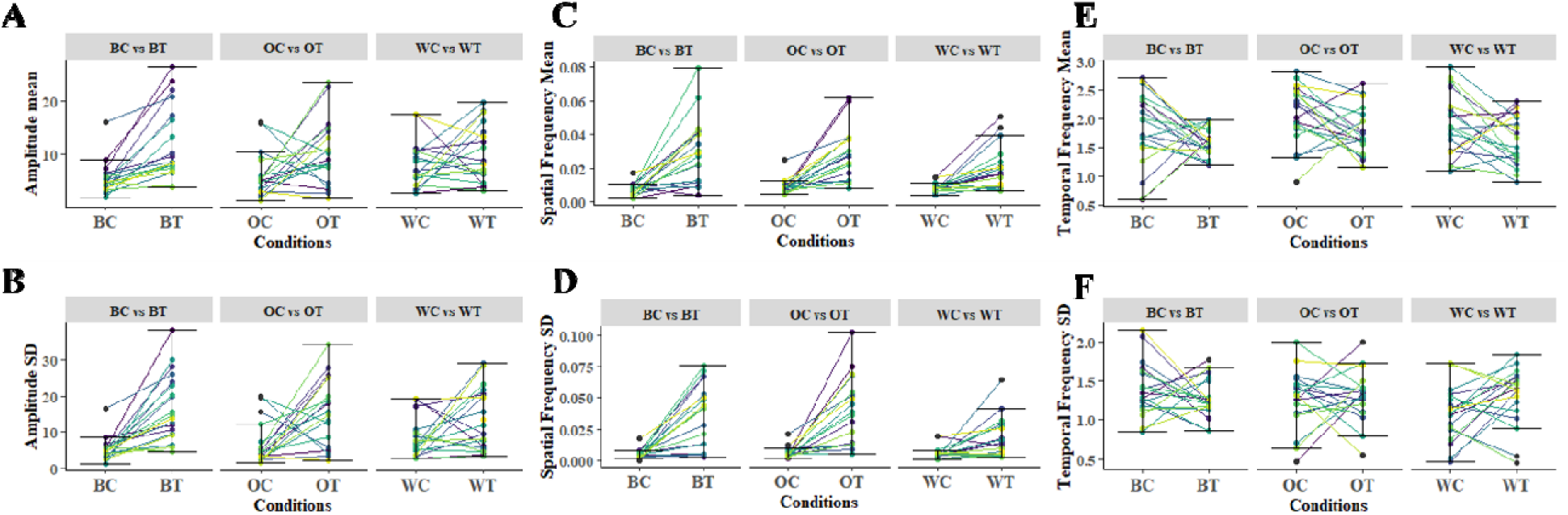
Path oscillations. The plots show the effects of colour change on Amplitude mean, Spatial frequency mean, Temporal frequency mean, Amplitude SD, Spatial frequency SD, and Temporal frequency SD from the control to the test conditions. BC, WC and OC represent Black control, White control, Orange control, whereas BT, WT and OT represent Black, White, Orange tests, respectively. Units are cm for amplitude, cycles per cm for spatial frequency, and cycles per second for temporal frequency.

In order to understand the phase relation between path oscillations and orientation oscillations, we plotted the starts of orientation cycles on to cycles of path oscillations, defining as 0° both the start of turning left and the start of turning right (Figure 7). The angular distributions confirmed that overall, starts of head turns were scattered and uniformly distributed on the path-oscillation cycle, suggesting little by way of coupling between the two kinds of cycles. Nevertheless, a few ants in some conditions show a preponderance at 0°, hinting at some coupling of the oscillations.

**Figure 7.**
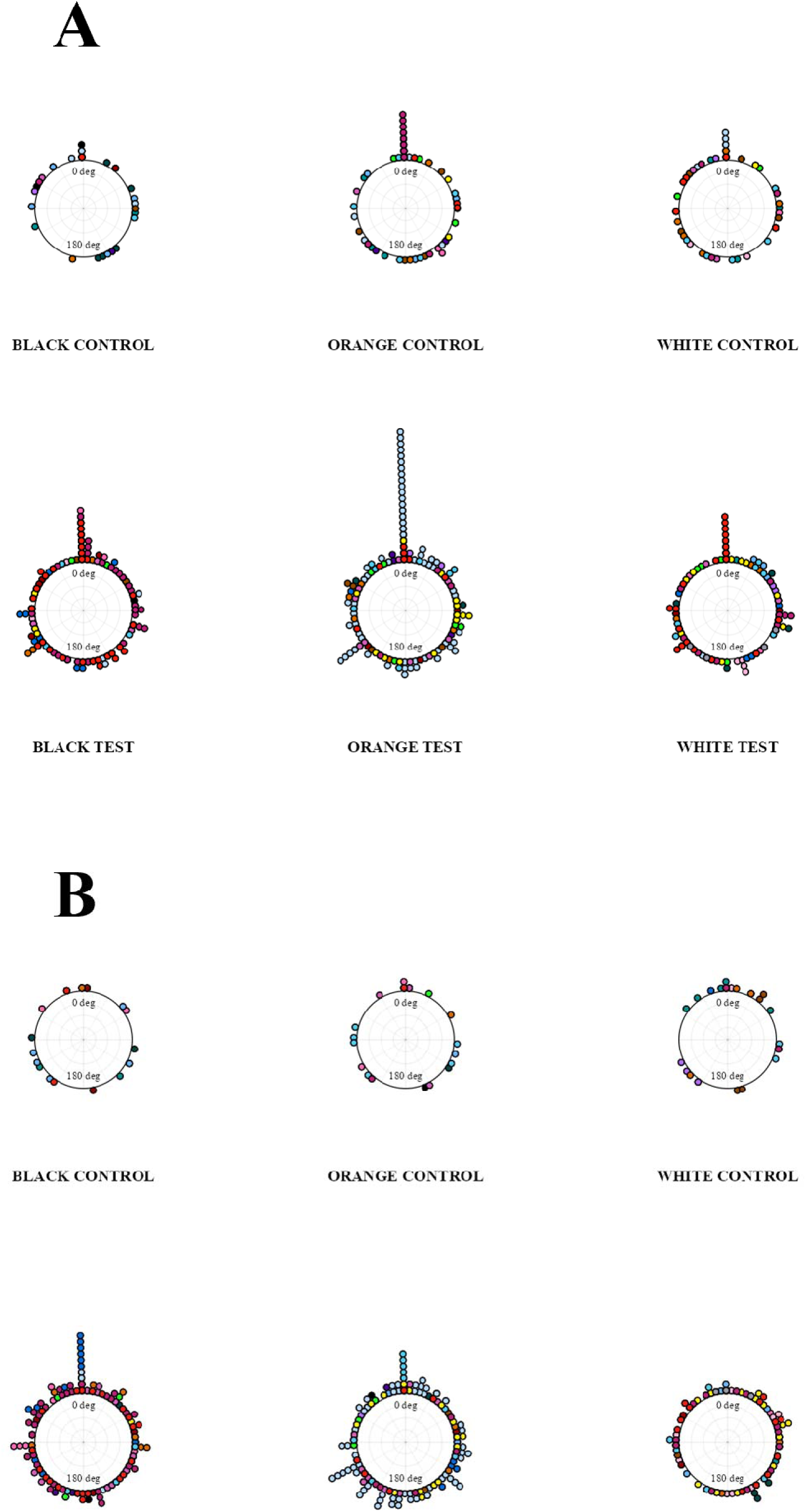
Phase relationships of control and test conditions (Left to right) and (Right to left). Circular histograms show the starts of left turns in orientation oscillations relative to the starts of left turns in path oscillations (0 deg) (A) and the starts of right turns in orientation oscillations relative to the starts of right turns in path oscillations (0 deg) (B). Angles increase clockwise on the graphs.

To help to find a psychophysical signature associated with the colour change, we calculated the effect sizes for all variables measured (Table 1). For measures of oscillations, the effect sizes vary, but for speed, straightness measures, duration of travel, and scans, Cohen’s d is consistently above 1. As 0.8 is considered a large effect, all these measures show large or sometimes very large effect sizes. This group could furnish a psychophysical signature of change detection.

## DISCUSSION

Changing the colour of the boards prominently placed before the feeder had large effects on the travel of desert ants towards their usual feeder. The ants slowed down their travel, meandered more on multiple measures of meander, scanned more and longer, and their path and orientation oscillations changed as well. The spatial frequency of path oscillations increased, consistently across all the colour-change conditions, while it was the amplitude of orientation oscillations that increased consistently. This suite of changes allows us to formulate a ‘signature’ package for conducting visual psychophysics in the natural habitat of ants and gives insights into the organisation of oscillatory behaviour in the course of navigation, two topics that we discuss in more detail.

### Visual psychophysics in the natural habitat

We used changes that we thought would be obvious to the ants. The purpose was not to demonstrate that the ants can detect such a change, but to come up with a signature suite of measurements that can be used in a program of visual psychophysics. The focus of psychophysics is on the visual stimulus changes, more particularly in this case, the changes that the species can detect, rather than on the navigational behaviour. It is not necessary to measure and compute all the variables that we assessed; a subset will do. For psychophysics, the ease of measurement and computation is a desideratum, as is robustness across different visual changes. We suggest not measuring oscillatory characteristics, which require much extraction and computation, with some of them not robust across the conditions of this study.

The speed, straightness or meander measures, and the scanning provide enough of a signature, being robust and easy to obtain.

We recommend Straightness (*D*/*L*), number of scanning bouts, and duration of the walk on video as a suite. For these measures, only a single point on the ant has to be tracked and extracted, and all these measures require minimal computations, easily done on a spreadsheet. The effect sizes (Cohen’s d) of all measures across all conditions are > 1. It is possible that the first Principal Component of these three variables in a Principal Components analysis (Figure 8) might serve even better. Indeed, on such an analysis, we see only a single case of an ant in the ‘wrong’ direction.

**Figure 8.**
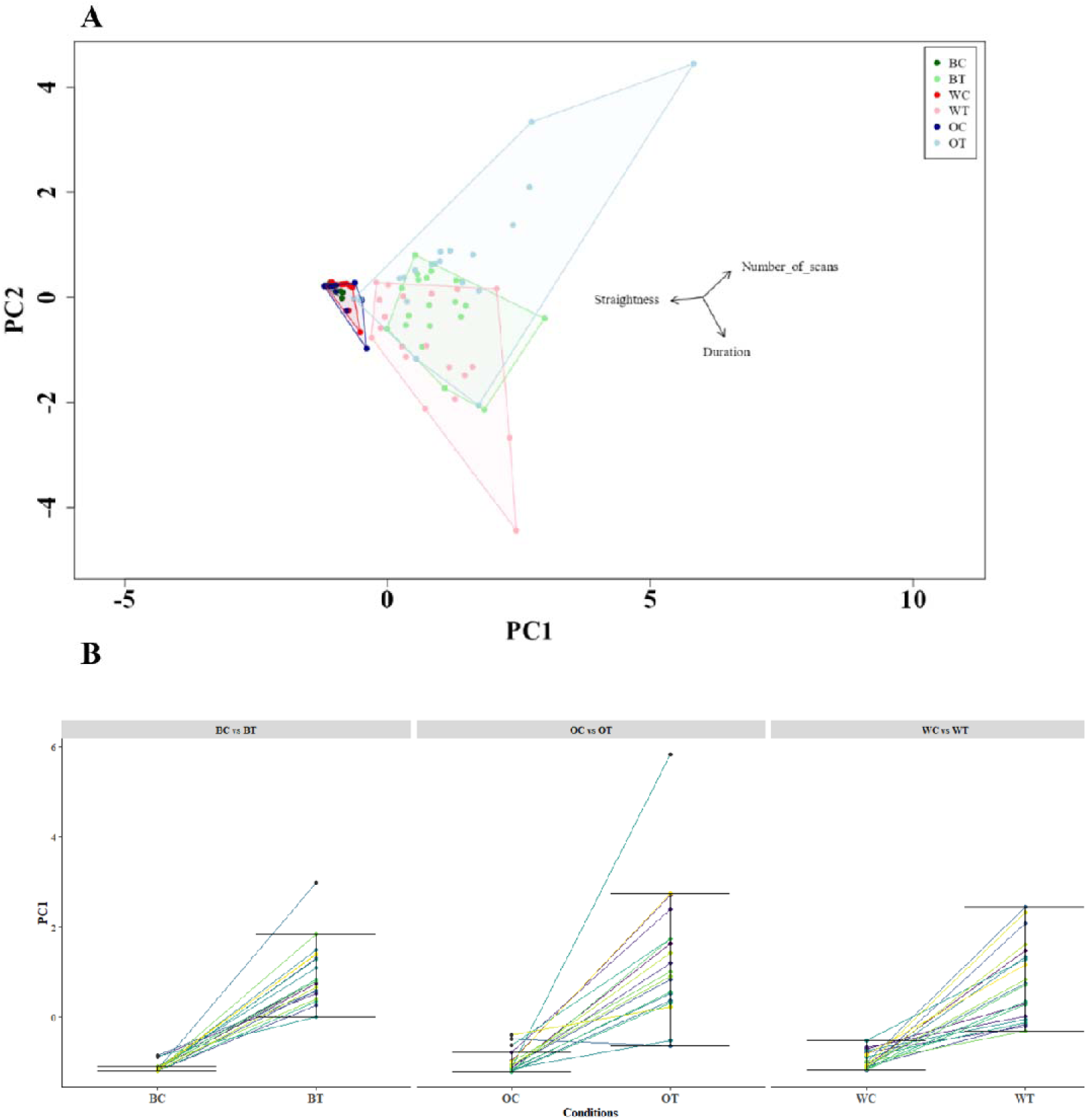
Principal Component Analysis (PCA) performed on Straightness, number of scans, and duration. A) The scatter plot shows the distribution of data points across the first two principal components (PC1 and PC2). B) The PC1 component. Means of the control and test conditions are shown for each individual. Data points in A) are colour-coded to represent different categories.

A psychophysical program could then use the signature to determine thresholds for detecting visual changes. Consider, for example, the visual width of vertical stripes that can be detected, a measure of spatial resolution in vision. One might train ants going to a feeder with uniformly grey boards at first. Replacing the grey boards with boards containing two vertical black stripes and two vertical white stripes should lead to detection, as indicated by the psychophysical signature. By gradually reducing the width of the stripes on the test boards with different groups of ants, at some point, the ants will not be able to detect any differences between the striped and the grey boards. Psychophysical titration allows researchers to estimate this kind of spatial resolution.

Such a psychophysical method has advantages and disadvantages. A major disadvantage is that a single animal is only tested once with the stimulus change. After one exposure, the changed stimulus condition is no longer new, so that a second test cannot be made—or, more precisely put, the second test would not be with a new stimulus. The psychophysical data need to be pooled across animals, with possible individual differences muddying the estimations. We have also demonstrated the proof of concept in only one species, so that further comparisons in other species would be good to have. Major advantages include the ease of measurement, which we have already argued for, the natural conditions of testing, which is in the habitat in which the ant would encounter visual stimuli in their lives, and the ease of training. Ants that find and take food from provisioned feeders do not need special training. For the red honey ants, if scientists set up a feeder and put a border encompassing the nest and feeder, the foragers will find it of their own accord.

### Oscillations in navigation

Our study also added details to how oscillations in ant navigation change with visual changes on the route, adding to initial studies on this topic (Clement et al., 2023; Murray et al., 2020). We can identify three kinds of oscillations: of the legs, of the head, and of the path. We did not examine leg oscillations (see Wilson, 1966, on insect gaits) because their analysis requires a higher frame rate. Head (orientation) and path oscillations are regular enough that straight- forward definitions have allowed us to describe them and analyse their properties. What we found was that with stimulus changes, head oscillations increase in amplitude, thus swinging wider from left to right and right to left, while path oscillations increased in spatial frequency. Both these changes are consistent with the function of sampling more of the visual world in different directions from different places along the route. The increased spatial frequency of path oscillations would have the concomitant effect of increasing measures of meander, all of which were significantly different between control and view-change conditions. The increase in scanning, both number of bouts and bout durations, and the slower speed of movement are also consistent with more visual sampling of the environment.

We thus found in this study that oscillations are adjusted in the services of navigation (Cheng, 2022, 2023). Oscillations enroute allow ants to sample the visual environment to some extent on navigational trips, even when conditions are normal and well-learned. Ants oscillate more, in the ways described above, when encountering visual changes. In other contexts, the meandering of paths increases when navigational journeys had not gone well. For example, when ants are captured by experimenters near their destination (nest in the studies) and placed back on their route, meandering increased and the ants scanned more (bull ants: Deeti et al., 2023; Lionetti et al., 2024; desert ants: Wystrach et al., 2019). When desert ants had fallen into a trap that delayed their journey home, meandering and scanning also increased on the subsequent journeys, in the area just before the trap (Wystrach et al., 2020). Oscillatory characteristics were not measured in these studies and further research on the ‘rewinding’ and trap manipulations could characterize the oscillations that we examined in this study. Given the prevalence of oscillations in locomotion, we would also recommend more studies of these basic units of action (Gallistel, 1980) in navigation (Cheng, 2022, 2023).

We examined phase the relationship between head orientations and path oscillations and failed to find consistent phase coupling across the conditions of the experiment. The lack of positive evidence, however, should not be taken to be strong support for the null hypothesis of no phase relationship. Because we only videotaped a small segment of each journey, each ant contributed few data points, especially in control conditions, when the ants moved quickly through the recording area. Still, some ants showed a preponderance at 0° on the circular histograms of phase relationships (Figure 7), meaning that the left or right turns of the head and path took place together in time. In future, it would be good to obtain data on head and path oscillations across an entire journey of individual ants.

## Conclusion

We have presented a proof of concept for a way to conduct visual psychophysical studies with ants in their natural habitat. The method relies on the ants’ reaction to detected visual changes. The procedure consists in getting ants familiar with a set of visual cues on their route to the feeder. After familiarization, the cues are changed. If the ants can detect the change, their travel characteristics will change. Looking at simple measures of path straightness (straight-line distance of path over path length), duration of travel through a recording area, and the number of scanning bouts, or their combination, provides a good signature of detection. We also found that visual changes on the route led to increased amplitude of head swings and increased spatial frequency of path oscillations.

## ACKNOWLEDGEMENTS

We acknowledge the traditional custodians of the land upon which this research was conducted, the Arrernte people. Their culture and customs have nurtured and sustained this land since the Dreamtime and continue to do so today. We pay our respects to their Elders past and present. We thank the Centre for Appropriate Technology at Alice Springs, Australia for letting us work on their property and providing some storage space, and the CSIRO Arid Zone Research at Alice Springs for administrative support. We are also thankful to Donald James McLean and Drew Allen for helping us with data analysis.

## Author contributions

Experimental design: SD. Data collection: SD and VL; Data analysis: SD. Writing: SD and KC.

## Ethics standards

Australia has no ethical regulations regarding work with insects. The study was non-invasive and no long-term aversive effects were found on the nests or on the individuals studied.

## Data availability

Supplementary videos, Excel file and R scripts are available at Open Science framework: https://osf.io/hj3cn/ and dryad: https://datadryad.org/stash/resources/317602/review

---

**Figure S1.**
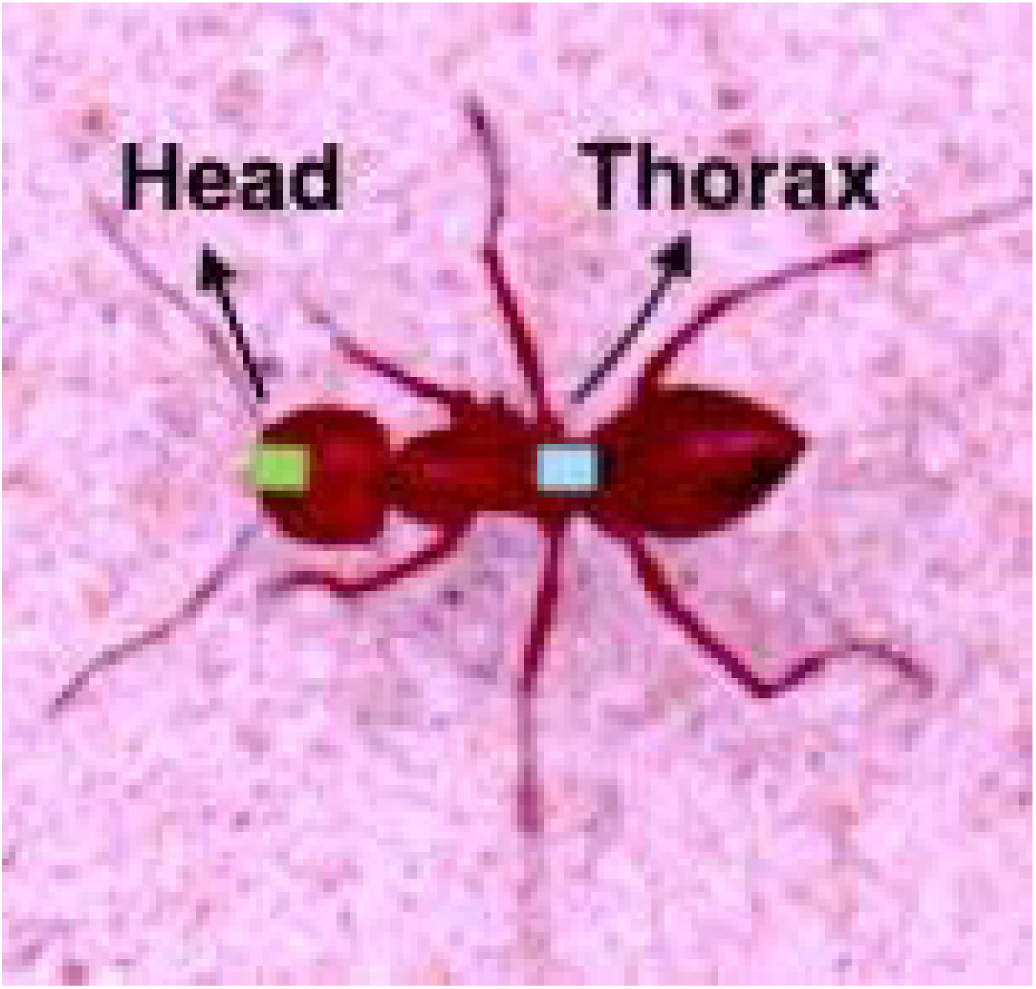
The annotated ants’ two body positions, used in data analysis: Front of the head and Mid of the thorax.

**Figure S2.**
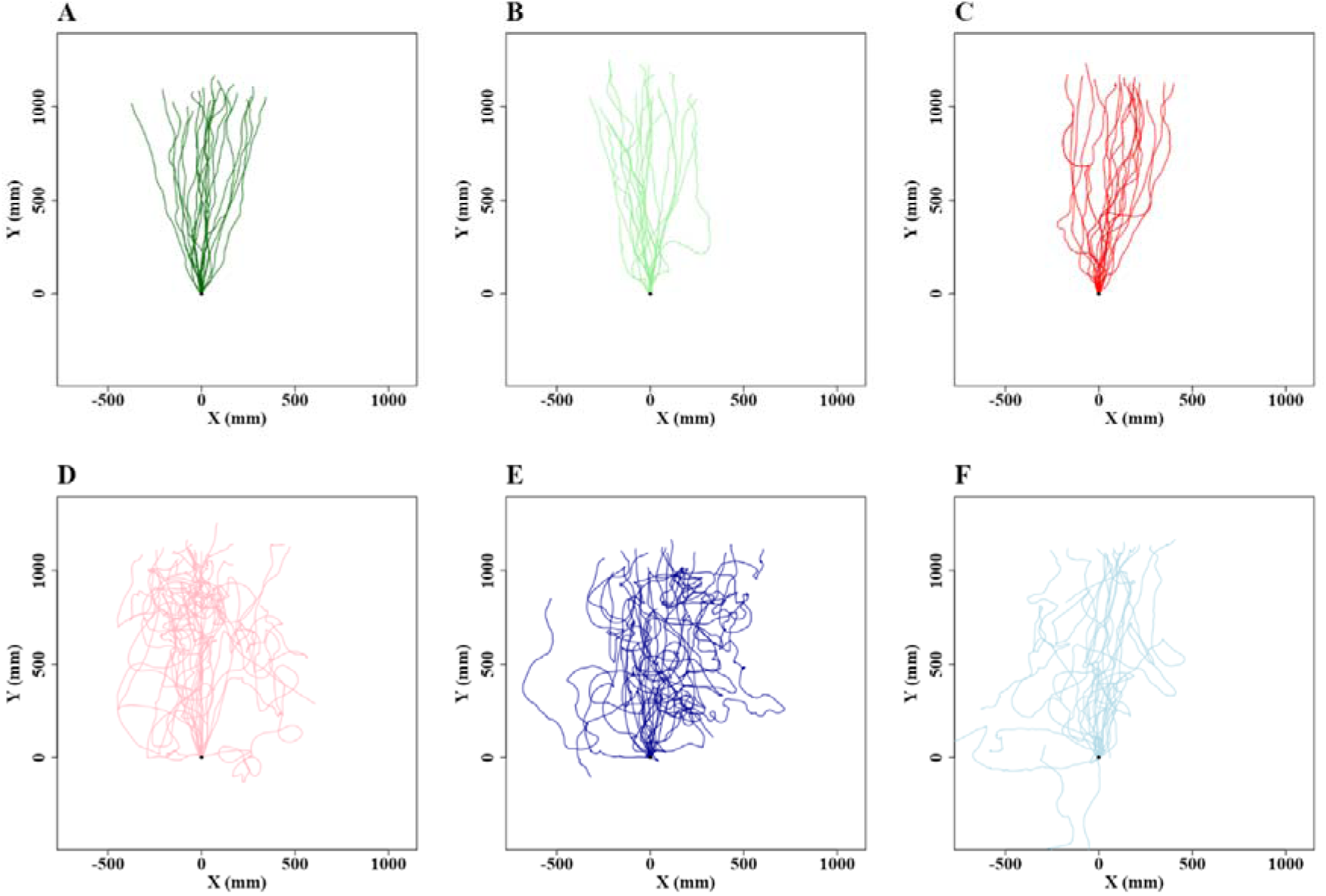
Trajectories of red honey ants in control and test conditions with various colours. (A) to (F) show the Control and Test paths of the individual desert ants during foraging. (A), (B), and (C) represent the control paths before the colour of the board changed. (D), (E) and (F) depict the test paths after the colour changes. (A), (D), Black to White; (B), (E), White to Orange; (C), (F), White to Black. Each trajectory plot represents the movement path of an individual ant. The y-axis points towards the goal. The coordinates (0,0) represent the starting frame of each recording.

