## Supplementary material for "A Method for Visual Psychophysics based on the Navigational Behaviour of Desert Ants (*Melophorus bagoti*)": supplimental data

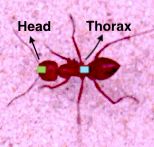

**Figure S1.** The annotated ants’ two body positions, used in data analysis: Front of the head and Mid of the thorax.

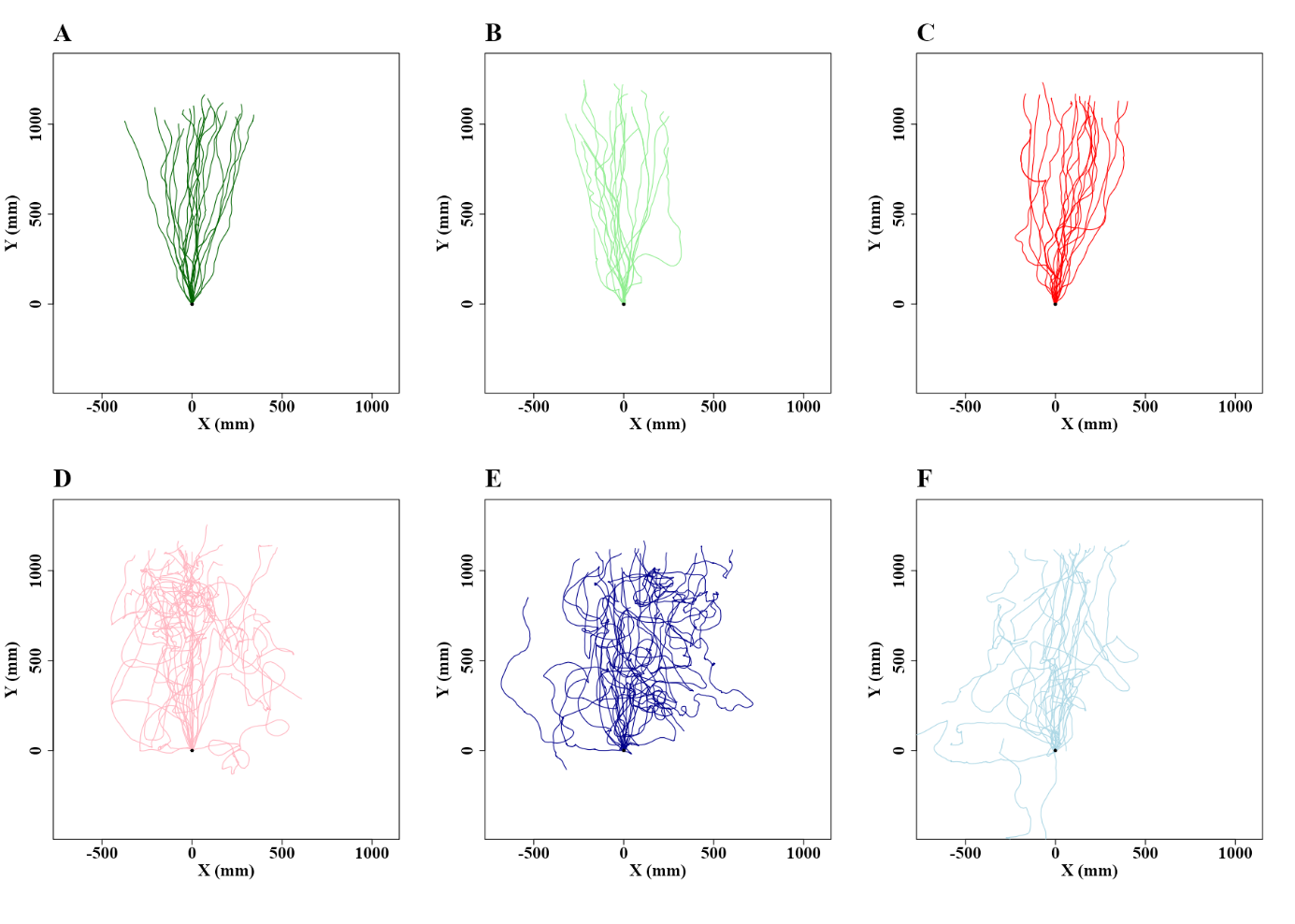

**Figure S2. Trajectories of red honey ants in control and test conditions with various colours.** (A) to (F) show the Control and Test paths of the individual desert ants during foraging. (A), (B), and (C) represent the control paths before the colour of the board changed. (D), (E) and (F) depict the test paths after the colour changes. (A), (D), Black to White; (B), (E), White to Orange; (C), (F), White to Black. Each trajectory plot represents the movement path of an individual ant. The y-axis points towards the goal. The coordinates (0,0) represent the starting frame of each recording.
